# ABC transporters support *Drosophila* intestinal stem and progenitor cells during cytotoxic challenge

**DOI:** 10.64898/2026.10.05.756803

**Authors:** Hannah Dayton, Jonathan DiRusso, Kristopher E. Kolbert, Olivia Williamson, Aiste Balciunaite, Edridge D’Souza, Kelly Becker, Elizaveta Hosage, Muneera Issa, Victoria Lu, Raghuvir Viswanatha, Shu Kondo, Norbert Perrimon, Michele Markstein

## Abstract

Adult stem cells can survive diverse insults, from ionizing radiation to toxic chemicals. In vertebrates, ABC transporter expression and dye-efflux capacity have been associated with stem-cell protection and multidrug resistance. How compound accumulation differs between adult stem cells and their differentiated daughters within an intact tissue remains poorly understood. Here, we show that adult *Drosophila* intestinal stem/progenitor cells maintain a distinct low-accumulation state for membrane-permeable compounds and that ABC transporters contribute to this phenotype and to the response to cytotoxic stress. Across nine membrane-permeable probes, stem/progenitor cells accumulated eight at lower levels than differentiated enterocytes. We selected tetramethylrhodamine methyl ester (TMRM), a membrane-permeable fluorescent probe that showed a strong difference in accumulation between stem/progenitor cells and enterocytes, as a quantitative readout for an RNAi screen of all 56 *Drosophila* ABC-family genes. The screen identified seven candidate transporters whose depletion increased the stem/progenitor-to-enterocyte TMRM fluorescence ratio. Further characterization of one of them, ABCB7, showed that *ABCB7*-depleted guts contained fewer mitotic cells during cytotoxic drug exposure. These results implicate ABC transporters in lower compound accumulation in *Drosophila* intestinal stem/progenitor cells and support a role for ABC transporters in their response to cytotoxic challenge.

## INTRODUCTION

ATP-binding cassette (ABC) transporters are transmembrane pumps found across organisms ranging from bacteria to humans^1^. Functional ABC transporters contain two transmembrane domains that form the substrate-translocation pathway and two nucleotide-binding domains that couple ATP binding and hydrolysis to transport; these domains may occur within a single polypeptide or be assembled through dimerization of half-transporters^2,3^. The substrates and direction of transport vary among individual transporters. In bacteria, ABC transporters mediate both import and export, whereas in eukaryotes many family members function as exporters, moving drugs, xenobiotics, lipids, and metabolites out of the cytosol or across intracellular membranes^4–6^. By reducing intracellular concentrations of drugs and other xenobiotics, plasma-membrane exporters can contribute to cellular protection^7,8^. Well-characterized examples include ABCB1/P-glycoprotein and ABCG2/BCRP, which export structurally diverse compounds and can reduce intracellular drug accumulation^8^. In vertebrates, elevated ABC transporter expression has been documented in several stem cell and cancer stem cell populations, where it can contribute to resistance to cytotoxic compounds, including some chemotherapy drugs^9–12^. This efflux capacity has also been used to identify stem-cell-enriched populations, most notably the hematopoietic “side population,” defined by exclusion of Hoechst 33342 and enriched for functional stem-cell activity^13^. ABCG2 contributes to this phenotype and is expressed in stem-cell populations from multiple tissues^14^.

This protective capacity may be particularly important in the intestinal epithelium, where dietary, microbial, and xenobiotic molecules are encountered continuously and long-lived stem cells must persist while generating differentiated daughter cells specialized for absorption and digestion^15^. Differentiated intestinal epithelial cells express several ABC transporters, including ABCB1/P-glycoprotein, ABCC2/MRP2, and ABCG2/BCRP, which contribute to the intestinal barrier by limiting xenobiotic absorption^16,17^. These studies identify ABC transporter activity as an important determinant of chemical exposure in the mature intestinal epithelium, but much less is known about how stem/progenitor cells handle the same compounds. A Hoechst-low side population enriched for putative intestinal epithelial stem cells has been isolated from mouse jejunum, suggesting that a high-efflux state may also occur in intestinal stem cells^18^. ABC transporter-mediated protection is also present in the stem-cell niche: Wnt-producing intestinal myofibroblasts are resistant to several xenobiotics, and loss of *Mrp1/Abcc1* increases their drug sensitivity^19^.

The adult *Drosophila* midgut provides a genetically tractable system in which to examine these relationships between epithelial cell identity, chemical exposure, and stem/progenitor-cell behavior. It is maintained by intestinal stem cells (ISCs), which self-renew and generate absorptive enterocytes (ECs) and secretory enteroendocrine (EE) cells^20,21^. Enterocytes arise through an enteroblast (EB) progenitor state, and ISCs and EBs express *escargot* (*esg*), allowing genetic manipulation of stem/progenitor cells separately from differentiated ECs^22^. ECs make up most of the intestinal epithelium and form a polarized absorptive layer facing the gut lumen, whereas ISCs and EBs occupy a basal position adjacent to the basement membrane^21,23^. In response to epithelial damage, multiple signaling pathways, including JAK–STAT, EGFR/MAPK, JNK, and Hippo/Yorkie, regulate ISC proliferation and epithelial regeneration^24–27^.

ABC transporter function is also established as a component of chemical protection in *Drosophila*. At the blood–brain barrier, the Mdr65 transporter limits xenobiotic entry and protects against cytotoxic compounds, and expression of human ABCB1/MDR1 can restore transport function in flies lacking *Mdr65*^28^. Loss of *Mdr49*, *Mdr50*, or *Mdr65* alters insecticide susceptibility, with effects that depend on the transporter and compound tested^29^. Tissue-specific knockdown studies have identified additional contributions to pesticide responses, while localization studies show distinct distributions of P-glycoprotein family members within the midgut^30^. Notably, Mdr50, one of the transporters implicated in these studies, also emerged as a candidate in our screen. Previous work showed that several chemotherapeutic drugs stimulate proliferation of intestinal stem cells through JAK–STAT signaling^31^. Damage-induced signaling helps explain why these cells proliferate, but whether intestinal stem/progenitor cells differ from their differentiated daughters in compound accumulation has not been established. It is also unclear which ABC transporters contribute to this phenotype and to stem/progenitor-cell responses during cytotoxic stress.

Here, we examined compound accumulation and ABC transporter function in the adult *Drosophila* intestine. We found that stem/progenitor cells accumulated most membrane-permeable fluorescent dyes at lower levels than neighboring enterocytes. Because tetramethylrhodamine methyl ester (TMRM) is a membrane-permeable fluorescent probe that produced a strong and quantifiable difference between these cell populations, we used it as the readout for an RNAi screen of all 56 *Drosophila* ABC-family genes. We reasoned that if an ABC transporter contributed to maintaining low TMRM accumulation in stem/progenitor cells, its knockdown should increase TMRM fluorescence in these cells relative to neighboring enterocytes. The screen identified seven candidate genes whose knockdown increased TMRM accumulation in stem/progenitor cells relative to neighboring enterocytes. Reporter analysis prioritized *CG32091* and *ABCB7* for follow-up, and functional analysis supported a role for *ABCB7* in the stem/progenitor response to cytotoxic challenge.

## RESULTS

We examined cytotoxic stress responses in the adult *Drosophila* intestinal epithelium, which is directly exposed to ingested compounds. Like the human intestine, the fly intestine is composed of a single-layered epithelial tube that is regionalized along its length for distinct digestive and absorptive functions. The *Drosophila* midgut performs digestive and absorptive functions analogous to the mammalian small intestine^15^. We focused on its posterior R5 region, one of the physiologically distinct regions of the adult midgut^32,33^ (**Figure 1a**). Large polyploid enterocytes, the major differentiated cell type in this region, form a brush border epithelium at their apical ends^20^ which, like in mammals^34^, is specialized for carrying out absorptive functions (**Figure 1a**). Following EC loss, intestinal stem cells^20,21^ divide to replenish the epithelium. In the EC lineage, enteroblasts serve as progenitors that differentiate into ECs^22–24,35^ (**Figure 1b**). ISCs and progenitors can be distinguished from ECs by their expression of *escargot* (*esg*)^20^, which is required to maintain stem/progenitor identity and prevent differentiation in intestinal stem/progenitor cells^22,35^. The ECs can be distinguished by their expression of the brush border cytoskeletal protein Myosin 1A (Myo1A)^36^. In the present work, we used *esg* (*esg-GAL4*)^20^ and *Myo1A* (*Myo1A-GAL4*)^24^ to mark and manipulate the stem/progenitor cells and ECs, respectively (**Figure 1b**).

**Figure 1.**
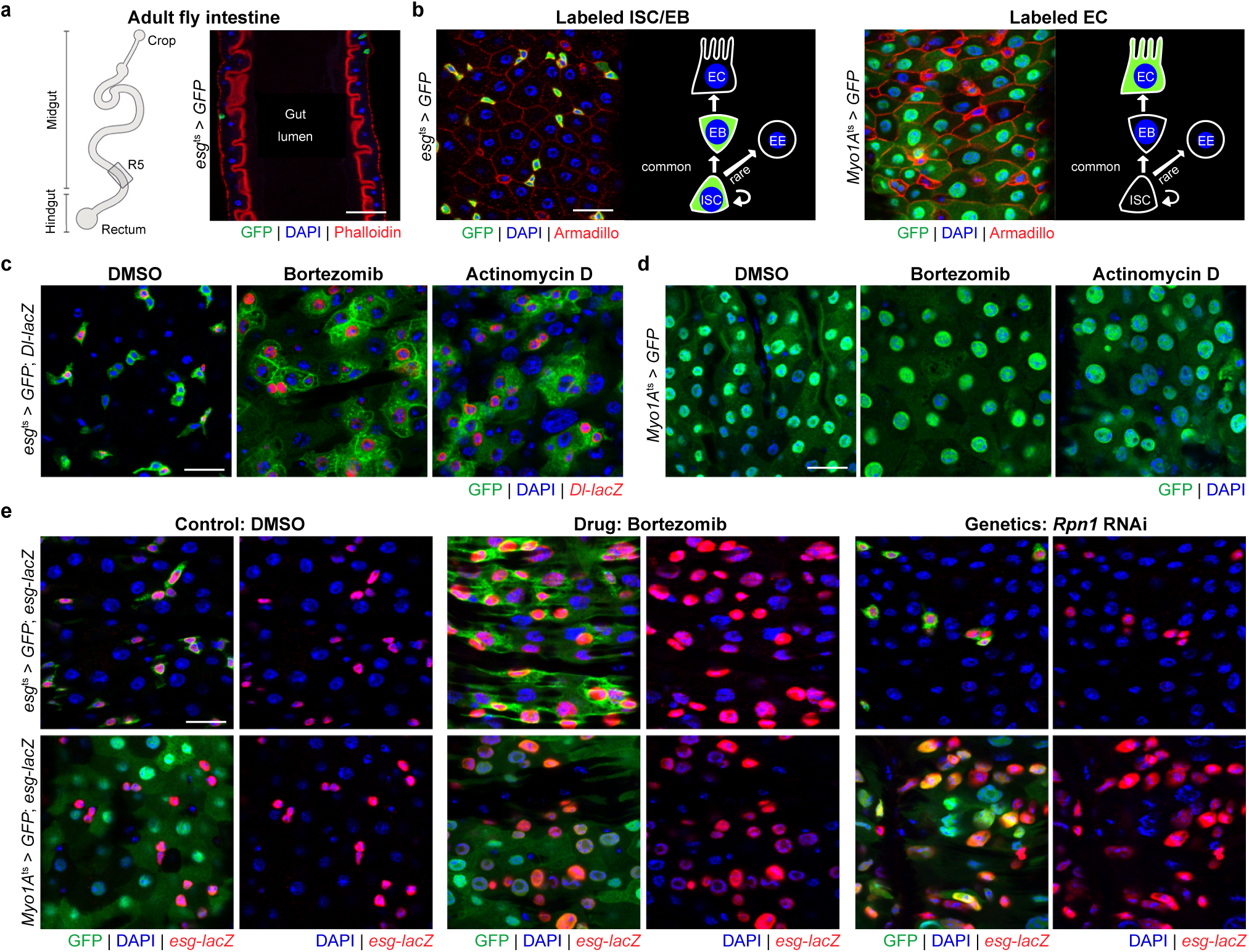
Cell-type-dependent responses to cytotoxic drugs and disruption of essential cellular processes. (**a**) Schematic and confocal cross-section showing the R5 region of the adult *Drosophila* intestine. Phalloidin staining of F-actin shows the rich brush border of ECs lining the gut lumen and muscle lining the outside of the gut. Stem/progenitor cells, marked in green, are positioned basally, abutting the surrounding muscle. Nuclei are stained with DAPI (blue). (**b**) Intestinal stem cell lineage and surface views of the gut epithelium showing stem and progenitor cells marked by *esg-GAL4* driving GFP and ECs marked by *Myo1A-GAL4* driving GFP. (**c**) Effect of drugs on ISCs (*Dl-lacZ*+, red) and progenitors (*Dl-lacZ*−) within the *esg*+ population (green). (**d**) Effect of drugs on ECs (green). (**e**) Effect of the proteasome-targeting drug bortezomib and RNAi against *Rpn1* or *mCherry* control driven by *esg-GAL4* and by *Myo1A-GAL4*. *esg*+ cells are marked with *esg-lacZ* in red in all panels in e. Scale bars, 15 µm.

To investigate the differential effect of cytotoxic drugs on ISCs vs. ECs, we used the stem cell-specific reporter *Delta-lacZ* (*Dl-lacZ*)^24,37^ to track stem cells in flies treated with two structurally and mechanistically distinct chemotherapy drugs, bortezomib, a proteasome inhibitor, and actinomycin D, a transcription inhibitor. In previous work, we have shown that these drugs induce ECs to express the JAK–STAT cytokine Upd3, which in turn induces *esg*+ cells to proliferate^31^. However, we did not distinguish the stem cells from progenitors nor assess the effects on EC survival. Following drug treatment, *Dl-lacZ*+ ISCs remained readily detectable and appeared more abundant in the imaged regions (**Figure 1c**). Conversely, drug-treated intestines appeared to contain fewer ECs within the imaged regions, and the remaining ECs appeared larger, with enlarged nuclei (**Figure 1d**). This morphology is consistent with previous observations of compensatory EC enlargement following genetically induced EC death^24^. These observations suggest that ECs and stem/progenitor cells respond differently to the same cytotoxic exposures.

We next used the GAL4-UAS TARGET system^38,39^ to disrupt essential cellular processes selectively in stem/progenitor cells or ECs. To compare genetic disruption of proteasome function with bortezomib treatment, we used *esg-GAL4* or *Myo1A-GAL4* to express RNAi against *Rpn1*, which encodes a proteasome regulatory subunit^40^. *Rpn1* depletion in ECs increased the abundance of *esg*+ cells (marked with *esg-lacZ*), resembling the response to bortezomib feeding, whereas depletion in stem/progenitor cells reduced the *esg*+ population (**Figure 1e**). Further, EC disruption alone expanded the *esg*+ population, consistent with previous findings that damaged ECs stimulate ISC proliferation^26,41^. We observed a similar cell-type-dependent pattern following knockdown of *α-Tubulin at 84B* (*αTub84B*): direct disruption of microtubule function in stem/progenitor cells reduced the *esg*+ population, whereas *αTub84B* knockdown in ECs increased *esg*+ cell number (**Supplementary Figure 1**). These contrasting cell-type-dependent responses prompted us to examine whether stem/progenitor cells and ECs differ in intracellular compound accumulation.

We reasoned that if stem/progenitor cells maintain lower intracellular levels of ingested small molecules, this difference should be detectable using membrane-permeable fluorescent probes. To test this prediction across chemically distinct compounds, we fed flies a panel of nine membrane-permeable fluorescent dyes and compared their fluorescence in *esg*+ stem/progenitor cells and ECs. The panel included two rhodamine derivatives, TMRM and rhodamine 123, as well as seven SYTO Red dyes ranging in molecular mass from approximately 400 to 650 Da.

Because ISCs/EBs lie basally beneath the enterocyte layer (**Figure 1a**), lower fluorescence could also result if ingested probes had reduced access to these cells. We therefore also tested 40-kDa tetramethylrhodamine-conjugated dextran (TMR-dextran), a large, membrane-impermeable tracer that does not freely diffuse across plasma membranes and is too large to be an expected substrate for ABC-transporter-mediated efflux. TMR-dextran served as an accessibility control to determine whether ingested fluorescent material could reach the basally located stem/progenitor cells.

Across the panel, eight of nine membrane-permeable probes accumulated at lower levels in *esg*+ stem/progenitor cells than in differentiated ECs (**Figure 2a,b**). **Pattern I** probes, including SYTO 60 and TMR-dextran, labeled both *esg*+ ISCs/EBs and surrounding enterocytes. SYTO 60 is a small, membrane-permeable dye that labeled both stem/progenitor cells and enterocytes. In contrast, TMR-dextran is a large, membrane-impermeable tracer, so its detection in stem/progenitor cells indicates that ingested fluorescent material can reach these cells despite their basal position. These **Pattern I** observations show that stem/progenitor cells are accessible to ingested fluorescent material and can also accumulate at least some membrane-permeable probes.

**Figure 2.**
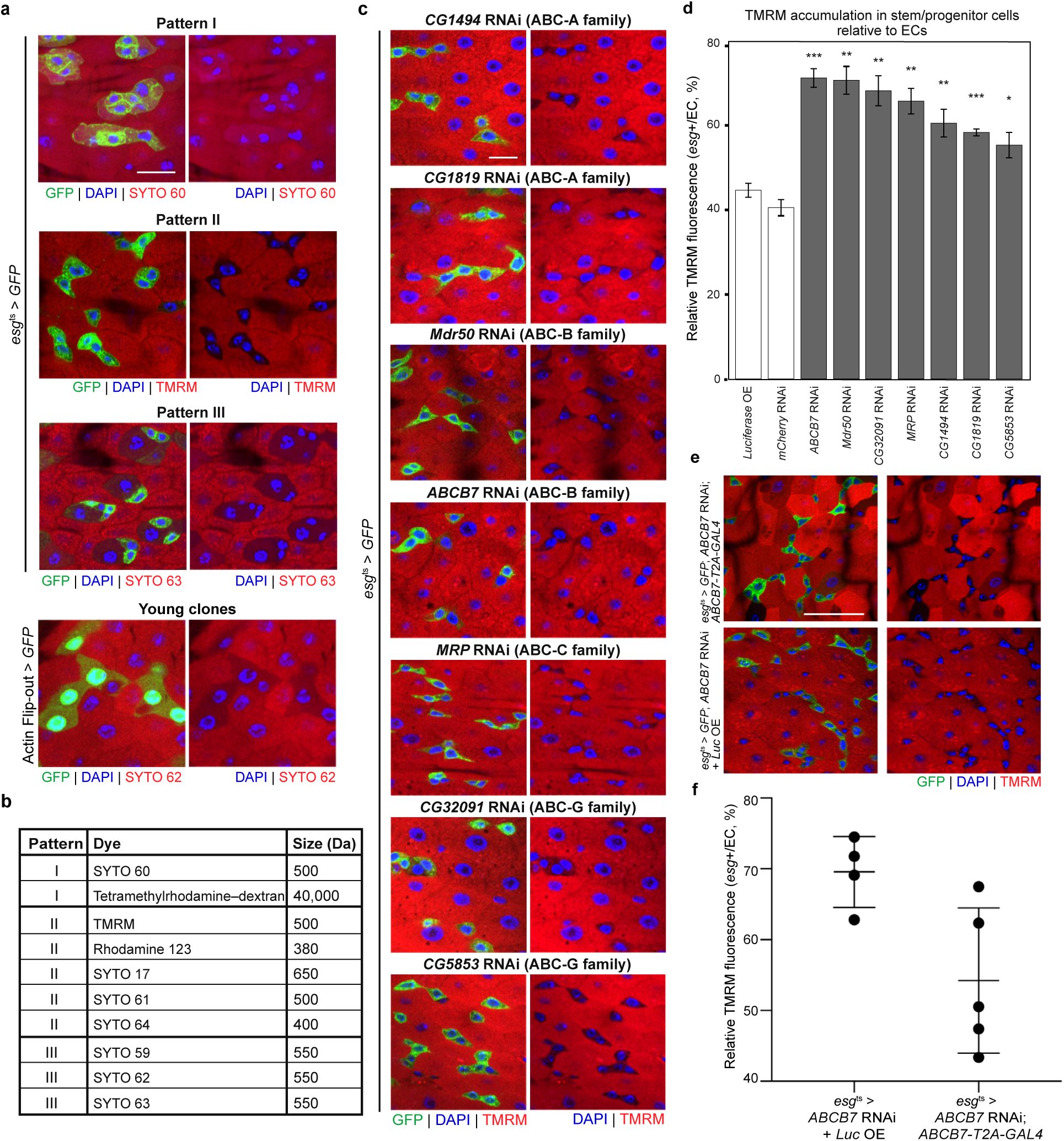
ABC transporter knockdown alters relative TMRM accumulation in intestinal stem/progenitor cells. (**a**) Representative dye-accumulation patterns following two days of feeding. GFP marks stem/progenitor cells or Actin flip-out clones (green), dyes are shown in red, and DAPI marks nuclei (blue). Patterns I–III and SYTO 62 accumulation in young clones are shown. GFP is omitted from paired right-hand images. Scale bar, 25 µm. (**b**) Classification and approximate molecular mass (Da) of the probes tested. (**c**) Representative images of TMRM accumulation following knockdown of seven candidate ABC transporters. GFP marks *esg*+ cells (green), TMRM is red, and DAPI is blue. GFP is omitted from paired right-hand images. Scale bar, 10 µm. (**d**) Relative TMRM accumulation, calculated as the ratio of mean fluorescence intensity per pixel in *esg*+ cells to neighboring ECs within each gut (%). At least four guts were analyzed per genotype. Error bars indicate SD. Two-tailed Welch’s *t*-tests: *** *p* < 0.001, ** *p* < 0.01, * *p* < 0.05. (**e**) Representative images following *ABCB7* RNAi with *UAS-luciferase* as a non-rescue control or the RNAi-resistant *ABCB7-T2A-GAL4* allele. GFP is green, TMRM is red, and DAPI is blue. Scale bar, 30 µm. (**f**) Quantification of relative TMRM accumulation as in d. Each point represents one gut; horizontal bars indicate the mean and error bars indicate SD. *n* = 4 control and *n* = 5 rescue; *p* = 0.026, two-tailed Welch’s *t*-test. The rescue allele used in e–f carries the *βTub56D* 3′ UTR and lacks the endogenous *ABCB7* 3′ UTR RNAi target sequence.

The remaining membrane-permeable probes accumulated at lower levels in *esg*+ cells than in differentiated ECs and fell into two additional patterns. **Pattern II** dyes, including TMRM, accumulated at low levels in *esg*+ ISCs/EBs but strongly in neighboring ECs. **Pattern III** dyes showed low accumulation in *esg*+ ISCs/EBs and in some nearby ECs. For SYTO 62, Actin-Flp-out lineage tracing showed that young lineage-marked ECs also retained low fluorescence (**Figure 2a**), indicating that the low-accumulation state can persist beyond loss of stem/progenitor identity. These results show that lower compound accumulation is common across chemically distinct membrane-permeable probes in *esg*+ stem/progenitor cells, while the pattern of accumulation in ECs differs among probes.

We selected TMRM for follow-up because it produced bright fluorescence and a clear Pattern II phenotype, with markedly lower signal in *esg*+ stem/progenitor cells than in neighboring ECs, making the difference between the two cell populations readily quantifiable. TMRM is a small, membrane-permeable cationic dye whose cellular fluorescence depends on intracellular dye accumulation but is also influenced by mitochondrial membrane potential^42^. Because changes in mitochondrial physiology can alter TMRM fluorescence independently of transport, we used relative TMRM fluorescence as the screen readout rather than interpreting it as a direct measure of ABC-mediated efflux. Because many ABC transporters can reduce intracellular compound accumulation through active export, and because membrane-permeable dye assays have been used to identify efflux-capable vertebrate stem cells (e.g., the classic ‘side population’ assay^13^), we performed an RNAi screen against all 56 ABC genes encoded in the *Drosophila* genome^43^ and measured TMRM accumulation in *esg*+ cells relative to neighboring ECs (**Figure 2c,d**). For each gut, we quantified the ratio of mean TMRM fluorescence per pixel in *esg*+ stem/progenitor cells to that in neighboring ECs, providing an internally normalized measure of the difference in dye accumulation between the two cell populations. The screen identified seven candidate ABC transporter genes whose knockdown increased the *esg*+/EC TMRM fluorescence ratio relative to controls (**Figure 2c,d; Supplementary Table I**). These candidates span four fly ABC subfamilies: *CG1494* and *CG1819* (ABC-A), *Mdr50* and *ABCB7* (ABC-B), *MRP* (ABC-C), and *CG32091* and *CG5853* (ABC-G). These seven candidates provided a basis for examining transporter expression and function in intestinal stem/progenitor cells.

We next examined the expression patterns of the candidate transporters. We inserted a *T2A-GAL4* reporter^44–47^ immediately upstream of the endogenous stop codon in five candidate genes (**Figure 3a**) and obtained a MiMIC-*GFP* reporter^48^ for one of the others (**Figure 3b**). Of the six candidates examined, two showed reporter patterns consistent with stem/progenitor cells: *CG32091* and *ABCB7* (**Figure 3c**). Reporter signal was not detected in the stem/progenitor cells for the other four candidates under the conditions tested. *CG32091* reporter expression resembled the distribution of *esg*+ stem/progenitor cells under both vehicle and drug-treatment conditions (**Figure 3c**). In contrast, *ABCB7* reporter expression was not detectable under vehicle conditions but became detectable in *esg*+ stem/progenitor cells following treatment with bortezomib or actinomycin D. *ABCB7* transcript abundance in dissected R5 tissue increased following actinomycin D treatment with both primer pairs (**Supplementary Figure 2**). Because actinomycin D also increases the *esg*+ fraction, this bulk increase is consistent with either higher expression per cell or a larger population of *ABCB7*-expressing cells. *ABCB7* was an unexpected candidate because it is best characterized as a mitochondrial transporter. We therefore examined the subcellular distribution of ABCB7-GFP in *Drosophila* S2R+ cells. Transiently expressed ABCB7-GFP showed prominent peripheral signal with limited overlap with the mitochondrial marker ATP5A (**Supplementary Figure 3**).

**Figure 3.**
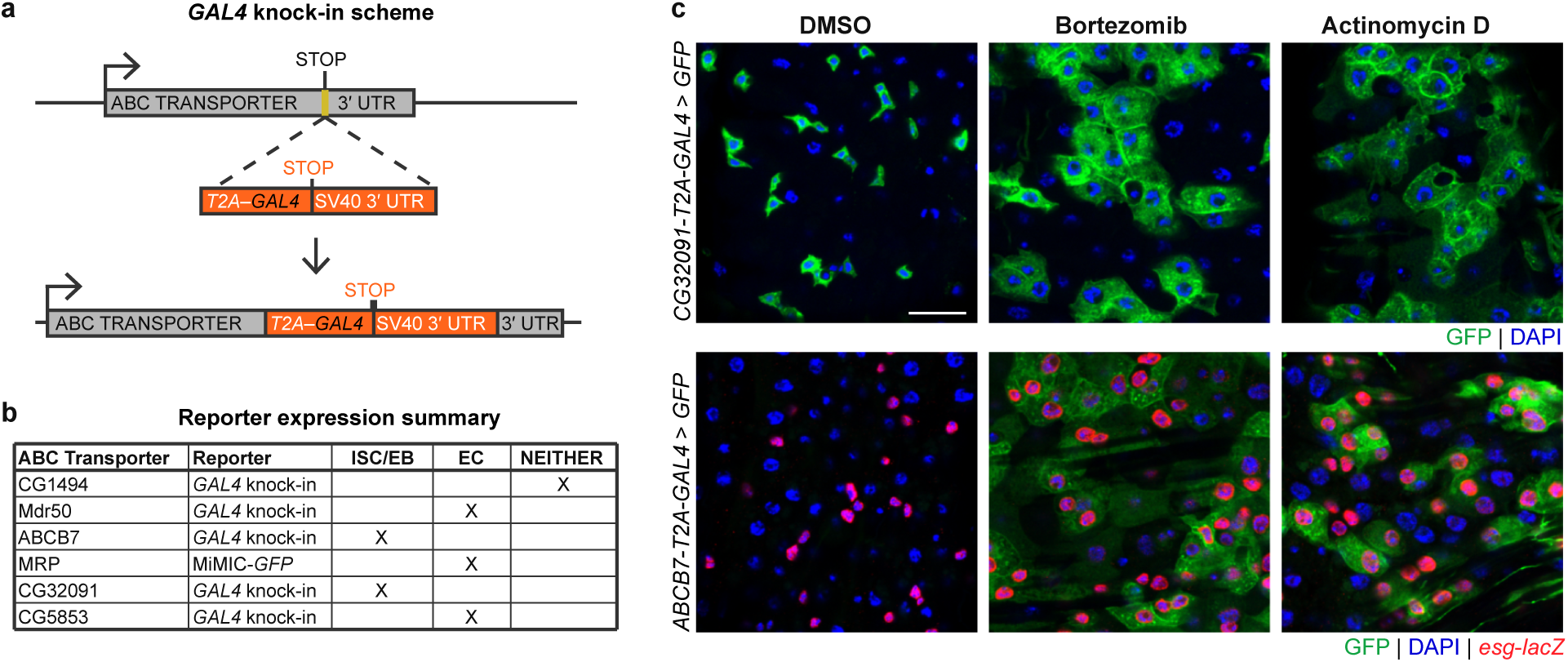
Expression of candidate ABC transporters. (**a**) Scheme for generating *T2A-GAL4* knock-in reporters by insertion of *T2A-GAL4* immediately upstream of the endogenous stop codon and in frame with the final coding exon. The inserted *SV40* 3′ UTR terminates the transcript upstream of the endogenous 3′ UTR, creating a transcriptional reporter for each gene. (**b**) Summary of expression patterns of ABC transporters identified in the RNAi screen whose knockdown increased the *esg*+/EC TMRM fluorescence ratio. Two reporters showed patterns associated with stem/progenitor cells; the remaining four were not detected in these cells under the conditions tested. (**c**) Expression patterns of *CG32091* and *ABCB7* reporters under vehicle (DMSO) and following bortezomib or actinomycin D treatment. GFP marks reporter activity (green), and DAPI marks nuclei (blue). In the *ABCB7* images, *esg-lacZ* independently marks stem/progenitor cells (red). Scale bar, 15 µm. *The ABCB7-T2A-GAL4* reporter allele used in panel c carries the *SV40* 3′ UTR and is distinct from the RNAi-resistant allele used in Figure 2 and Supplementary Figure 4.

To test whether the *ABCB7* RNAi phenotypes reflect on-target knockdown, we took advantage of the fact that the TRiP hairpin HMC03142 targets the endogenous *ABCB7* 3′ UTR. In the *ABCB7-T2A-GAL4* rescue allele, the *T2A-GAL4* cassette is followed by the *βTub56D* 3′ UTR, so the endogenous *ABCB7* 3′ UTR, including the HMC03142 target sequence, is absent from the resulting transcript (**Supplementary Figure 4a**). We first used a developmental eclosion assay as an organismal test of hairpin specificity. In this assay, *esg-GAL4*-driven *ABCB7* RNAi was induced during development, and the fraction of RNAi-expressing pupae that successfully eclosed was scored in the presence or absence of the RNAi-resistant allele. Whereas RNAi in the adult gut experiments was induced only after eclosion, progeny in the eclosion assay were shifted to 29 °C during development. *ABCB7* knockdown strongly reduced eclosion, with 0/3 and 3/19 pupae eclosing in two independent experiments. Introduction of the RNAi-resistant allele restored eclosion to 10/11 and 34/38, respectively (**Supplementary Figure 4b**). In the adult gut, the same RNAi-resistant allele also partially rescued the elevated stem/progenitor-to-EC TMRM fluorescence ratio caused by *ABCB7* knockdown, reducing it from 69.6 ± 5.0% to 54.2 ± 10.2% (mean ± SD; *n* = 4 and 5 guts; *p* = 0.026, two-tailed Welch’s *t*-test; **Figure 2e,f**). Because *esg* is expressed in imaginal, tracheal, and neural precursors during development, the eclosion phenotype cannot be attributed specifically to the intestine. Rescue of both developmental viability and the adult gut TMRM phenotype supports an on-target effect of the *ABCB7* hairpin.

Because the low-accumulation phenotype might change as stem/progenitor cells respond to cytotoxic stress, we asked whether it was retained during bortezomib-induced expansion. Following co-feeding with TMRM and bortezomib, the expanded *esg*+ population remained relatively TMRM-low compared with neighboring ECs, indicating that differential accumulation was maintained during cytotoxic challenge (**Supplementary Figure 5**).

TMRM is a reported P-glycoprotein substrate, and bortezomib has also been reported to be transported by P-glycoprotein, providing a precedent for shared transporter recognition of a fluorescent probe and a cytotoxic drug^49,50^. We therefore tested whether *ABCB7* and *CG32091*, identified through their effects on relative TMRM accumulation, also contribute to the stem/progenitor response to bortezomib and the mechanistically distinct drug actinomycin D. Depletion of either *CG32091* or *ABCB7* reduced the expansion of the *esg*+ population following bortezomib or actinomycin D treatment (**Figure 4a**). We focused subsequent quantitative analyses on *ABCB7*. Following bortezomib or actinomycin D treatment, *ABCB7*-depleted guts contained fewer *esg*+ cells per R5 field than controls (**Figure 4a; Supplementary Figure 6**). *ABCB7*-depleted guts had fewer phospho-histone H3-positive (pH3+) mitotic cells than controls during bortezomib and actinomycin D exposure (**Figure 4c, Supplementary Figure 7**). *ABCB7*-depleted intestines retained an increase in *esg*+ cell number following dextran sulfate sodium (DSS) exposure (**Figure 4b,d**), indicating that depletion does not abolish this expansion response.

**Figure 4.**
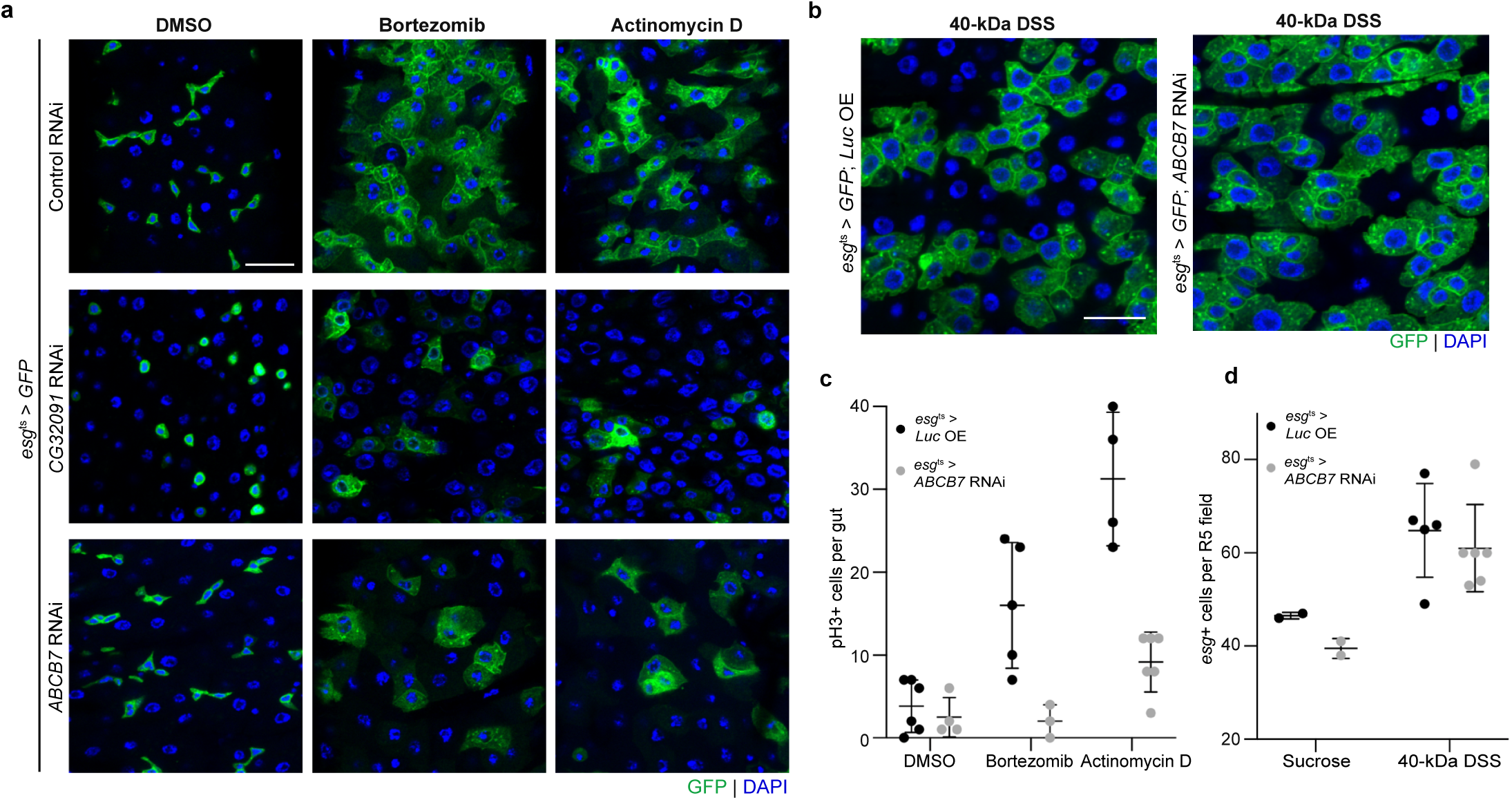
Effects of *ABCB7* and *CG32091* depletion on the intestinal stem/progenitor response to cytotoxic challenge. (**a**) Representative images of *esg-GAL4*-driven GFP-positive stem/progenitor cells (green) following control, *CG32091*, or *ABCB7* RNAi and treatment with DMSO, bortezomib, or actinomycin D. Scale bar, 15 µm. (**b**) Representative images of stem/progenitor cells in control and *ABCB7* RNAi intestines following treatment with 3% DSS in 4% sucrose. Scale bar, 15 µm. (**c**) Quantification of pH3+ cells in control (*esg*^ts^ > *Luc OE*) and *ABCB7* RNAi (*esg^ts^* > *ABCB7* RNAi) intestines following DMSO, bortezomib, or actinomycin D treatment. Each point represents one gut. Horizontal bars indicate the mean, and error bars indicate SD. DMSO: *n* = 6 control and *n* = 4 *ABCB7* RNAi; bortezomib: *n* = 5 control and *n* = 3 *ABCB7* RNAi; actinomycin D: *n* = 4 control and *n* = 6 *ABCB7* RNAi. (**d**) Quantification of *esg*+ cells per imaged field in R5 following sucrose or DSS treatment, with one field analyzed per gut. Each point represents one gut; horizontal bars indicate the mean and error bars indicate SD.

## DISCUSSION

We show that *Drosophila* intestinal stem/progenitor cells maintain a lower compound-accumulation state than their differentiated daughters within the same epithelium. This difference extended across several membrane-permeable probes tested, and depletion of multiple ABC-family genes altered relative TMRM accumulation. These results identify differential compound accumulation as a feature that distinguishes the stem/progenitor cells from neighboring enterocytes and implicate ABC transporters in this distinction. We also found that *ABCB7* contributes to the stem/progenitor response during exposure to cytotoxic drugs, suggesting a role for some ABC transporters in compound accumulation and cellular behavior under chemical challenge.

A useful precedent for interpreting this lower-accumulation phenotype comes from vertebrate side-population cells, which exclude fluorescent dyes such as Hoechst 33342 through ABC transporter-dependent mechanisms and are enriched for stem-cell activity^13,14^. Unlike vertebrate side-population cells, which are defined by their ability to exclude Hoechst dye, the *Drosophila* stem/progenitor cells in our experiments were identified independently by *esg* expression rather than by their dye-accumulation phenotype. Reduced fluorescence across eight of nine membrane-permeable probes therefore provides a phenotypic parallel with vertebrate dye-efflux populations, and the identification of multiple ABC-family genes in our screen strengthens this connection. ABC transporter-mediated efflux is likely to contribute to the lower compound accumulation observed in *Drosophila* stem/progenitor cells, although our experiments measure steady-state accumulation rather than efflux directly. Consistent with a protective role for ABC transporters during chemical stress, loss of *Abcg2* in mice increases hematopoietic-cell sensitivity to mitoxantrone^51^. Similarly, *ABCB5* depletion increases sensitivity to 5-fluorouracil in colorectal cancer^52^.

The low-accumulation phenotype was not restricted to the stem/progenitor state. Lineage tracing with SYTO 62 showed that young lineage-marked ECs retained low fluorescence after loss of the stem/progenitor state, indicating that the low-accumulation phenotype can persist during early EC differentiation. Changes in transporter expression could contribute to these differences, while changes in intracellular binding, organelle composition, membrane properties, or metabolic activity could also alter probe retention. Because the probes differ in their chemical properties and intracellular behavior, their distinct accumulation patterns may reflect different aspects of EC maturation.

The lower-accumulation state may be particularly relevant during cytotoxic drug exposure. The opposing responses to oral cytotoxic drugs and direct genetic disruption of essential cellular processes in *esg*+ cells indicate that stem/progenitor cells are not intrinsically insensitive to cellular damage; instead, their response to drug treatment likely reflects both their own cellular state and regenerative signals from damaged neighboring ECs. During bortezomib co-feeding, the expanded *esg*+ population remained relatively TMRM-low, showing that the low-accumulation phenotype is maintained during cytotoxic challenge.

The seven candidates span four ABC subfamilies and include both multidrug-transporter family members and transporters with other known cellular functions. Of the candidates examined with reporters, *CG32091* and *ABCB7* showed reporter patterns associated with stem/progenitor cells. *ABCB7* was particularly interesting because its reporter was not detectable in stem/progenitor cells at baseline but became detectable following bortezomib or actinomycin D treatment. *ABCB7* transcript abundance also increased following actinomycin D treatment, consistent with the reporter response observed under cytotoxic stress. Partial rescue with an RNAi-resistant *ABCB7* allele supports an on-target contribution of ABCB7 to the relative TMRM fluorescence phenotype. ABCB7 is best known as a mitochondrial transporter involved in cytosolic iron-sulfur protein maturation^53^. Because TMRM fluorescence is sensitive to mitochondrial physiology, the effect of *ABCB7* depletion on relative TMRM fluorescence could reflect altered mitochondrial function rather than direct transport of TMRM. To examine whether ABCB7 localizes exclusively to mitochondria, we assessed the distribution of transiently expressed ABCB7-GFP in S2R+ cells. ABCB7-GFP showed prominent peripheral signal with limited overlap with the mitochondrial marker ATP5A, suggesting that ABCB7 can localize outside mitochondria in this context. Its endogenous localization in intestinal stem/progenitor cells remains to be determined. ABCB7 has also been linked to proliferative capacity in mammalian cells: conditional loss of *Abcb7* in mice disrupts hematopoiesis^54^, and ABCB7-deficient B cells show impaired proliferation and replication stress^55^. Our finding that *ABCB7*-depleted intestines contained fewer mitotic cells during cytotoxic drug exposure is consistent with a role for ABCB7 in supporting proliferative responses under these conditions.

The retained expansion response to DSS indicates that *ABCB7*-depleted stem/progenitor cells remain capable of expansion in response to intestinal damage. Whereas *ABCB7* depletion reduced the expansion of the *esg*+ population following bortezomib or actinomycin D treatment, *ABCB7*-depleted intestines retained an increase in *esg*+ cell number following DSS exposure. DSS is known to activate a regenerative JNK–Hh–JAK–STAT signaling program in the *Drosophila* midgut^56^, and the preserved *esg*+ expansion therefore indicates that *ABCB7*-depleted stem/progenitor cells retain the capacity to respond to this form of intestinal damage. Our findings suggest that ABC transporters contribute to differences in compound accumulation between intestinal stem/progenitor cells and enterocytes and support a role for ABC transporters in the stem/progenitor response to cytotoxic challenge.

## MATERIALS AND METHODS

### Fly Stocks

Stocks used were:

w; esg-GAL4, UAS-GFP, tub-GAL80^ts^

w; esg-GAL4, UAS-GFP, tub-GAL80^ts^/CyO; Dl-lacZ/TM6B, Tb

tub-GAL80^ts^/FM7; Myo1A-GAL4, UAS-GFP/CyO

y¹ w^67c23^; P{lacW}esg^k00606^/CyO (esg-lacZ, BDSC 10359)

*UAS*-RNAi and control lines were obtained from the Bloomington *Drosophila* Stock Center (BDSC) and Vienna *Drosophila* Resource Center (VDRC): *Rpn1* (BDSC 34348), *αTub84B* (BDSC 31389), *CG1494* (BDSC 57777), *Mdr50* (BDSC 35034), *ABCB7* (BDSC 51696), *MRP* (BDSC 38316), *CG1819* (BDSC 38356), *CG32091* (BDSC 57783), *CG5853* (BDSC 27668), *mCherry* RNAi (BDSC 35785), and *UAS-luciferase* (BDSC 35788). The complete list of ABC transporter RNAi lines used in the screen, including TRiP identifiers, vectors, and insertion sites, is provided in **Supplementary Table I**.

*T2A-GAL4* reporter alleles for *CG1494*, *Mdr50*, *ABCB7*, *CG32091*, and *CG5853* were generated by CRISPR-mediated knock-in immediately upstream of the endogenous stop codon, placing *T2A-GAL4* in frame with the final coding exon^44,45^. In the alleles used as transcriptional reporters, an SV40 3′ UTR terminates the reporter transcript upstream of the endogenous 3′ UTR. For reporter imaging, *T2A-GAL4* alleles were crossed to *UAS-GFP*, and GFP expression was used to visualize reporter activity. For the *ABCB7* reporter experiments shown in Figure 3, *esg-lacZ* was included as an independent marker of stem/progenitor cells.

Two independently generated *ABCB7-T2A-GAL4* alleles were used in this study. *ABCB7-T2A-GAL4*^SV^^40^ (*yw;; ABCB7-T2A-GAL4^SV^*^40^) carries the SV40 3′ UTR as described above and was used as the reporter allele in **Figure 3**. *ABCB7-T2A-GAL4^βTub56D^* (*yw;; ABCB7-T2A-GAL4^βTub56D^*) was generated independently; in this allele, the *T2A-GAL4* cassette is followed by the *βTub56D* 3′ UTR and a 3xP3-dsRed screening marker, such that the endogenous *ABCB7* 3′ UTR is no longer transcribed. Because the TRiP hairpin HMC03142 targets the endogenous *ABCB7* 3′ UTR, transcripts from this allele lack the RNAi target sequence. This allele was therefore used as an RNAi-resistant source of *ABCB7* in the rescue experiments shown in **Figure 2** and **Supplementary Figure 4**.

The *MRP* reporter was the MiMIC insertion Mi{MIC}MRP^MI00447^ (BDSC 31007), located in the *MRP* 5′ UTR intron.

Stocks were reared on standard Bloomington Formulation food (Genesee Scientific, 66-121). All RNAi crosses were maintained at 21 °C. Adults were aged for 3–5 days at 21 °C before experimental procedures.

### RNAi Experiments

We used the TARGET system with GAL80^ts^ for all RNAi experiments. Flies were reared at 21 °C and transferred to 29 °C at 3–5 days of age for 7 days to induce RNAi before assays were performed. The developmental induction protocol used for eclosion assays is described below under Eclosion Assays.

### Fluorescent Dye Assays

Dyes were mixed into 500 µL of low-melt fly food as previously described for drug-feeding assays^31^. The nine membrane-permeable dyes were diluted in DMSO and administered at a final concentration of 2.5 µM in the food. 40-kDa TMR-conjugated dextran was diluted in 1× PBS and administered at a final concentration of 5 µM. Flies were fed dyes for 2 days at 29 °C, after which guts were dissected, fixed, mounted, and imaged. Ten fluorescent probes were tested: TMRM (Thermo Fisher Scientific, T668), rhodamine 123 (Thermo Fisher Scientific, R302), TMR-conjugated dextran, 40,000 MW, neutral (Invitrogen, D1842), and the seven SYTO dyes included in the SYTO 17 and 59–64 kit (Invitrogen, S11340). Approximate molecular masses are from the Thermo Fisher Molecular Probes Handbook, Table 8.3.

### Low-Melt Fly Food

We used a previously developed fly food formulation^31^ optimized for incorporating drugs in low volumes. The formulation contains low-melt agarose and standard agarose in place of agar. Low-melt fly food was prepared in distilled water containing 2% (wt/vol) autoclaved yeast, 7% (vol/vol) corn syrup, and 1.5% (wt/vol) total agarose, consisting of 1 part standard agarose to 11 parts low-melt agarose. The food was stored at 4 °C, remelted by boiling, and cooled to 37 °C before drugs were mixed into the liquid food. The resulting food-drug mixtures solidified at approximately 30 °C to form soft gels suitable for fly feeding. In initial experiments, low-melt fly food was labeled with food-grade red coloring (McCormick) and mixed with a range of bioactive compounds. Red coloration was visible in the abdomen within 1 h of feeding, confirming ingestion of the low-melt fly food preparation.

### Midgut Dissections and Visualization

Flies were anesthetized with CO₂ prior to dissection. Midguts were dissected in 200 mM glutamic acid, 50 mM KCl, 40 mM MgSO₄, 4 mM Na₂HPO₄, and 2 mM MgCl₂, as previously described^21^. Guts were fixed for 25 min in 4% formaldehyde prepared in 100 mM glutamic acid, 24 mM KCl, 20 mM MgSO₄, 2 mM Na₂HPO₄, and 1 mM MgCl₂. Samples were imaged using a Zeiss LSM700 confocal microscope.

### Quantitative Dye-Accumulation Assay

Nikon NIS-Elements software was used to calculate the mean red fluorescence intensity per pixel in *esg*+ stem/progenitor cells and neighboring enterocytes from at least four guts per sample. For each gut, relative dye accumulation was calculated as the ratio of mean fluorescence intensity in *esg*+ stem/progenitor cells to that in neighboring ECs. Ratios were multiplied by 100 for presentation as percentages. Ratios were then averaged across all guts within each sample. Fold change was calculated by dividing the mean fluorescence-intensity ratio for each experimental condition by that of the corresponding control. *p*-values were calculated using a two-tailed Welch’s *t*-test. Genes were designated as candidates if knockdown increased the *esg*+/EC TMRM fluorescence ratio with an unadjusted *p* < 0.05 relative to control lines. No correction for multiple comparisons was applied across the 56 genes because the screen was used for candidate identification rather than definitive statistical inference; selected candidates were subsequently evaluated in independent expression and functional assays.

### Drug Feeding Assays

Bortezomib (LC Labs, Cat# B-1408) and actinomycin D (Enzo, Cat# BML-GR300) were prepared as 20 mM and 10 mM stocks in DMSO, respectively, and stored at −20 °C. Flies were placed on low-melt fly food prepared as previously described^31^, and working stocks were diluted into the food to final concentrations of 20 µM bortezomib and 5 µM actinomycin D. DMSO diluted in low-melt fly food was used as the vehicle control; DMSO did not exceed 1% (vol/vol) of the food in any condition. Drug treatments were administered for three days at 29 °C prior to dissection. For TMRM-bortezomib co-feeding experiments, flies were fed 2.5 µM TMRM and 20 µM bortezomib in low-melt fly food for 2 days at 29 °C prior to dissection.

### DSS Challenge

40-kDa dextran sulfate sodium (DSS; Thermo Fisher, J63606) was prepared as a 3% (wt/vol) solution in 4% (wt/vol) sucrose in water and administered on chromatography paper. The paper was replenished with 200 µL of solution twice daily, and flies were exposed to DSS for 3 days at 29 °C.

### Immunofluorescence and Microscopy

Tissues were prepared as previously described^23^. The following primary antibodies were used: mouse anti-Armadillo (DSHB, N2 7A1), 1:20; rabbit anti-β-galactosidase (Cappell/MP Biomedicals, #55976), 1:10,000; mouse anti-β-galactosidase (DSHB, 40-1a), 1:50; rabbit anti-phospho-histone H3 (Millipore, 06-570), 1:1,000; and mouse anti-ATP5A [15H4C4] (Abcam, ab14748), 1:1,000, for S2R+ cell staining only. F-actin was visualized with Alexa Fluor 647–phalloidin (Invitrogen, A22287), 1:100. The following secondary antibodies were used (Invitrogen): Alexa Fluor 555 donkey anti-mouse, Alexa Fluor 647 goat anti-rabbit, and Alexa Fluor 647 donkey anti-mouse, all at 1:1,000. Nuclei were stained with DAPI at 1:30,000. Adult females were anesthetized, decapitated, and dissected in PBS, and tissues were fixed for 20 min. Dissected intestines were blocked in 5% normal donkey serum for 1 h, incubated with primary antibodies overnight, incubated with secondary antibodies for 1.5 h, and stained with DAPI for 6 min. Samples were mounted in VECTASHIELD mounting medium (Vector Laboratories, H-1000-10) and imaged on Leica TCS SP2 and Zeiss LSM700 confocal microscopes using a 40× oil-immersion objective.

### Image Acquisition and Handling

Images were acquired on a Zeiss LSM700 using a 40× oil-immersion Apochromat objective with ZEN 2011 Black software. Scale bars were added using Fiji (ImageJ) and the LSM Toolbox plugin. Images used for presentation were subjected only to linear brightness and contrast adjustments applied uniformly across the entire image. Quantitative measurements were performed on unadjusted image data. Adobe Photoshop CC and Fiji were used for figure preparation.

### S2R+ cell culture, transfection, and immunofluorescence

*Drosophila* S2R+ cells were maintained at 25 °C in Schneider’s *Drosophila* medium supplemented with 10% fetal bovine serum and penicillin-streptomycin. For transient expression of ABCB7-GFP, 3 × 10⁶ cells were seeded in 6-well dishes containing Concanavalin A (ConA)-coated glass coverslips. Cells were transfected with 400 ng total DNA consisting of Actin-GAL4 and UAS-ABCB7-GFP plasmids at a 1:10 ratio using Effectene transfection reagent (Qiagen) according to the manufacturer’s instructions. Cells were allowed to express ABCB7-GFP for approximately 24 h.

For immunofluorescence, cells were fixed in 4% paraformaldehyde in PBS and permeabilized in TBS containing 0.2% Triton X-100. Cells were incubated for 1 h with mouse anti-ATP5A [15H4C4] (Abcam, ab14748; 1:1,000) in TBS containing 0.05% Triton X-100 and 5% BSA, followed by a 1-h incubation with Alexa Fluor 647-conjugated anti-mouse secondary antibody and Hoechst in the same buffer. GFP fluorescence was used to visualize ABCB7-GFP. Images were acquired by confocal microscopy.

### RT-qPCR

Flies were raised as previously described and fed actinomycin D as described in the drug-feeding assay at 29 °C in cohorts of 10. On day 3, females were dissected in 1× PBS, and guts were cut at the Malpighian tubules and at the bend between the R4 and R5 regions. Twenty-five to thirty females were dissected per condition for each biological replicate. Guts were placed in TRIzol on ice immediately after dissection. RNA was extracted using a standard chloroform extraction and alcohol precipitation protocol. RNA concentration and purity were assessed using a NanoDrop spectrophotometer. cDNA was synthesized using the iScript cDNA Synthesis Kit (Bio-Rad). qPCR was performed on an Agilent Mx3000P qPCR system using the iTaq qPCR kit (Bio-Rad). Each sample was analyzed in duplicate, and all experiments were performed with three biological replicates. *RpL32* (formerly *Rp49*) was used as the normalization control. Primer sequences were as follows:

*RpL32* F: 5′-CGGATGGATATGCTAAGCTGT-3′

*RpL32* R: 5′-GCGCTTGTTCGATCCGTA-3′

*ABCB7* pair 1 F: 5′-CACCATCGAGCACAATATCCACT-3′

*ABCB7* pair 1 R: 5′-GGTCAGAGCCTGCAGAATGTTATG-3′

*ABCB7* pair 2 F: 5′-CAGGTAGGCGAACGTGGAT-3′

*ABCB7* pair 2 R: 5′-GAGCCTGCAGAATGTTATGCTC-3′

*upd3* F: 5′-AAATTGAATGCCAGCAGTACG-3′

*upd3* R: 5′-CCTTGCTGTGCGTTTCGTTC-3′

### Cell Type Quantification

For the *esg*+ cell quantification shown, images of the R5 region were exported to Fiji and analyzed using the Cell Counter plugin. GFP-positive *esg*+ cells were counted within one imaged field per gut, with each gut treated as an independent biological replicate. Error bars represent SD.

### pH3 Counts

Phospho-histone H3 (pH3) counts were performed throughout the gut. Nuclei were scored as pH3+ only when they showed bright, discrete pH3 staining associated with condensed mitotic chromatin; diffuse nuclear staining was not counted. Both apical and basal focal planes were examined. Counts were carried out in an *esg-GAL4, UAS-GFP, tub-GAL80^ts^* background expressing either *UAS-luciferase* or *UAS-ABCB7* RNAi.

### Eclosion Assays

*w; esg-GAL4, UAS-GFP, tub-GAL80^ts^; UAS-ABCB7* RNAi/TM6B, *Tb* males were crossed to either homozygous *ABCB7-T2A-GAL4βTub56D* or *UAS-luciferase* females. Parents were allowed to mate and lay at 21 °C for 2 days, after which progeny were shifted to 29 °C. Larvae were collected daily as they left the food. Pupae were scored as Tubby or wild-type-shaped and monitored through the eclosion period. Wild-type-shaped pupae carried *UAS-ABCB7* RNAi, whereas Tubby pupae lacked the RNAi transgene and were used to confirm genotype assignment. Only animals that fully emerged from the pupal case were scored as eclosed. Animals that ruptured the operculum but failed to fully emerge from the pupal case and subsequently died were scored as failed eclosion events. Two independent experiments were performed per condition.

## ACKNOWLEDGEMENTS

We thank Kim Tremblay for critically reading the manuscript. HD, JD, and ED received support from the University of Massachusetts Amherst Commonwealth Honors College. ED received support from a Goldwater Scholarship. N.P. is an investigator of Howard Hughes Medical Institute. This article is subject to HHMI’s Immediate Access to Research policy, which requires that this article be made publicly available as initial and revised preprints deposited on a designated preprint server under a CC BY 4.0 license.

## DATA AVAILABILITY

All data are available in the main text, figures, supplementary information and supplementary data. Correspondence and requests for materials should be addressed to.

---

**Supplementary Figure 1.**
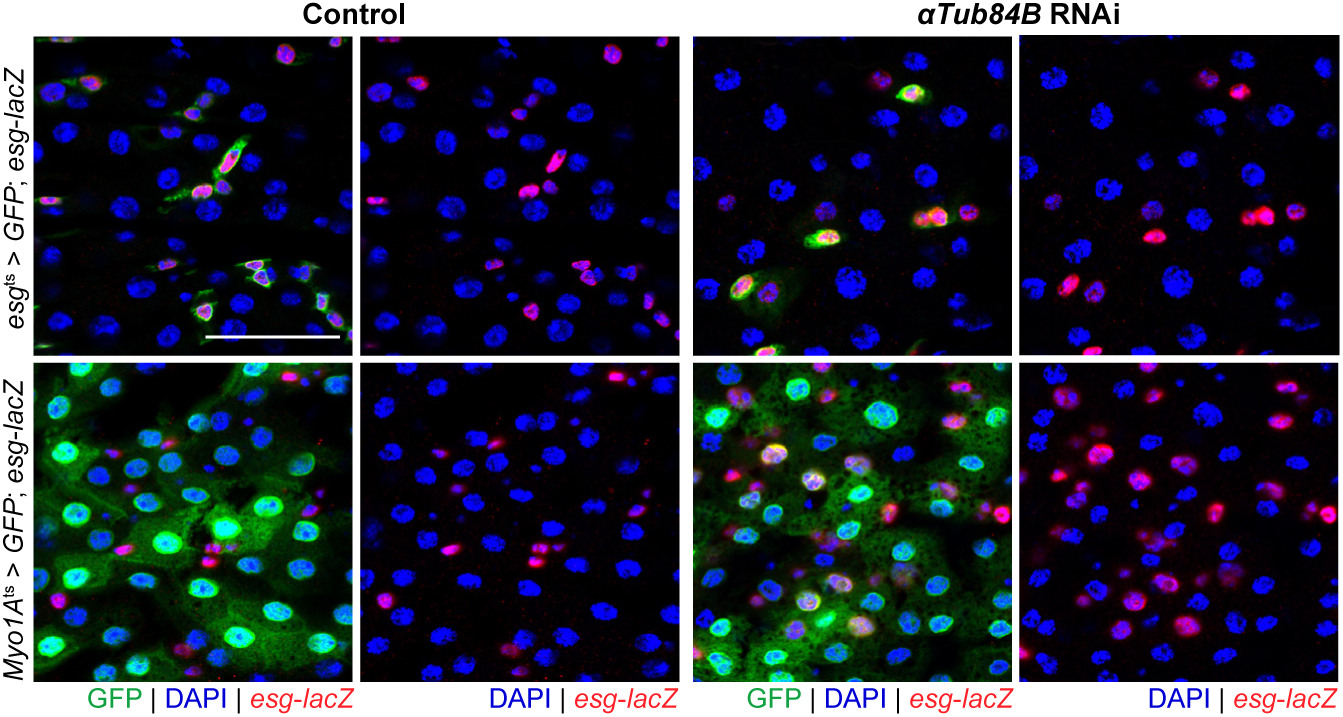
Cell-specific *αTub84B* knockdown produces opposing effects in stem/progenitor cells and enterocytes. Representative images of control *mCherry* RNAi and *αTub84B* RNAi in stem/progenitor cells or enterocytes. Top row: *esg-GAL4*-marked ISC/EB cells (green). Bottom row: *Myo1A-GAL4*-marked enterocytes (green). *esg-lacZ* marks stem/progenitor cells (red), and DAPI marks nuclei (blue). For each genotype, the left panels show all channels, and the right panels show *esg-lacZ* and DAPI only. *αTub84B* knockdown in stem/progenitor cells reduces the number of *esg*+ cells, whereas *αTub84B* knockdown in enterocytes increases the number of *esg*+ cells. Scale bar, 30 µm.

**Supplementary Figure 2.**
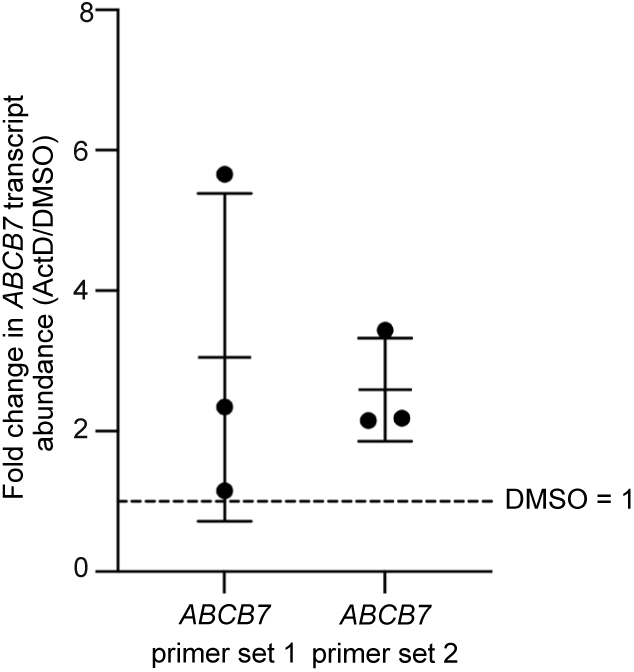
*ABCB7* transcript abundance following actinomycin D treatment. RT-qPCR analysis of dissected R5 gut tissue following 3 days of actinomycin D treatment. *ABCB7* transcript abundance is shown as fold change relative to the matched DMSO control for each biological replicate. Two *ABCB7* primer pairs were analyzed. Each point represents one biological replicate; *n* = 3 biological replicates per primer set, with 25–30 female guts pooled per replicate. Horizontal bars indicate the mean, and error bars indicate SD. The dashed line indicates no change relative to DMSO (fold change = 1).

**Supplementary Figure 3.**
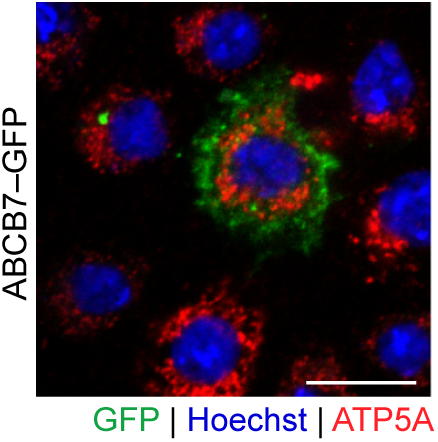
Subcellular distribution of ABCB7-GFP in *Drosophila* S2R+ cells. Representative image of transiently transfected S2R+ cells expressing ABCB7-GFP (green) and stained with anti-ATP5A to label mitochondria (red). Nuclei are stained with Hoechst (blue). Scale bar, 10 µm.

**Supplementary Figure 4.**
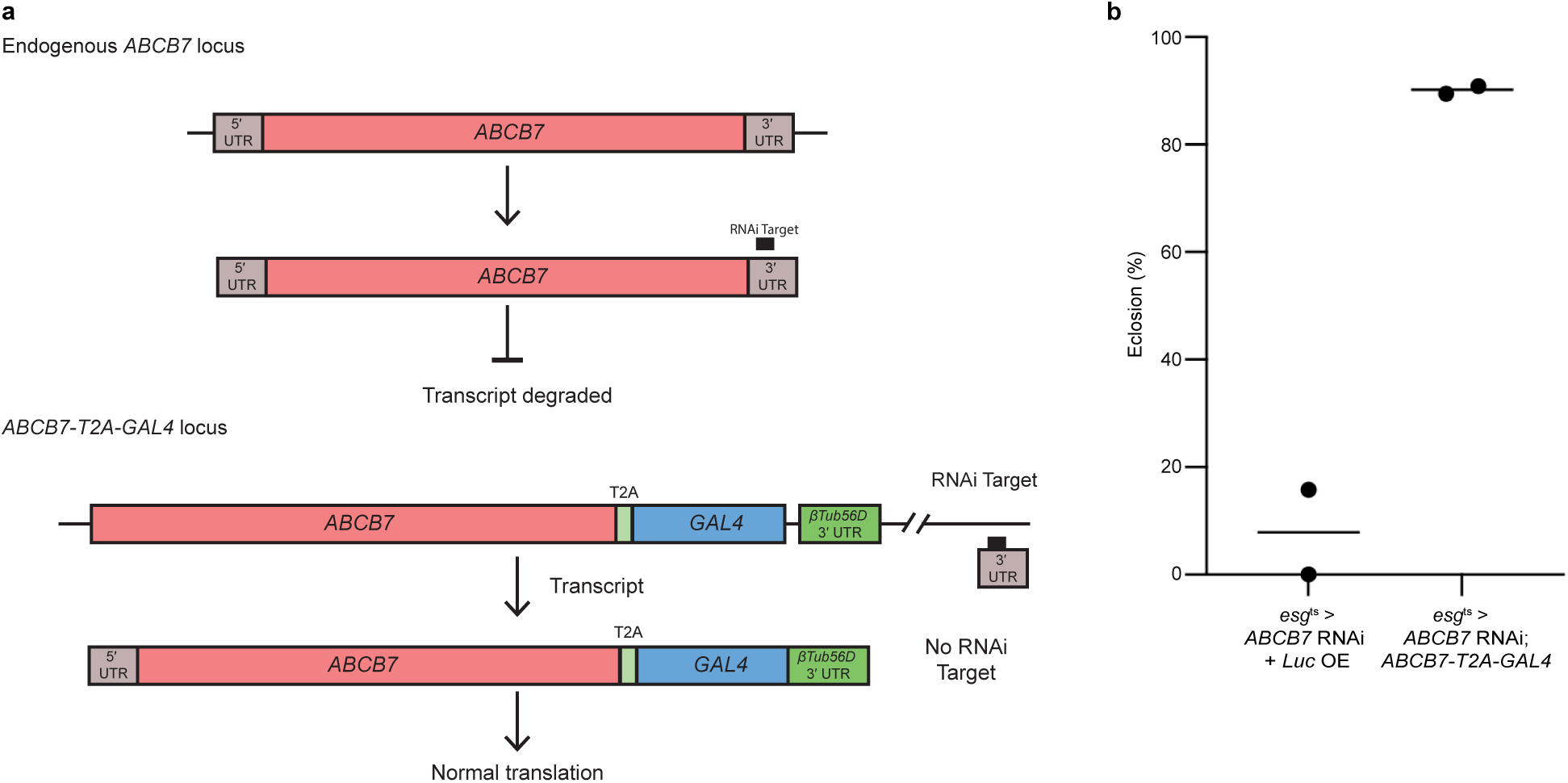
An RNAi-resistant *ABCB7* allele rescues phenotypes caused by *ABCB7* knockdown. (**a**) Schematic of the *ABCB7* RNAi-rescue strategy. The *ABCB7* RNAi construct targets the endogenous *ABCB7* 3′ UTR. In the *ABCB7-T2A-GAL4* allele, the *T2A-GAL4* cassette is followed by the *βTub56D* 3′ UTR, which terminates the transcript upstream of the endogenous *ABCB7* 3′ UTR and therefore excludes the RNAi target sequence. (**b**) Eclosion following *esg-GAL4*-driven *ABCB7* knockdown in the presence of either *UAS-luciferase* as a non-rescue control or the RNAi-resistant *ABCB7-T2A-GAL4* allele. Parents were mated at 21 °C for two days; eggs were shifted to 29 °C, and larvae were collected daily. Pupal morphology was used to distinguish experimental and balancer-containing progeny: wild-type-shaped pupae carried *UAS-ABCB7* RNAi, whereas Tubby pupae lacked the RNAi transgene. Eclosion rate was calculated as the percentage of wild-type-shaped pupae that fully emerged from the pupal case. Each point represents one independent experiment; across the two experiments, 22 experimental pupae were scored for *ABCB7* RNAi + *UAS-luciferase* and 49 experimental pupae were scored for *ABCB7* RNAi + *ABCB7-T2A-GAL4*. Horizontal bars indicate the mean (*n* = 2 independent experiments per condition). This *βTub56D* 3′ UTR-containing allele was used in the rescue experiments in panel b and Figure 2e–f.

**Supplementary Figure 5.**
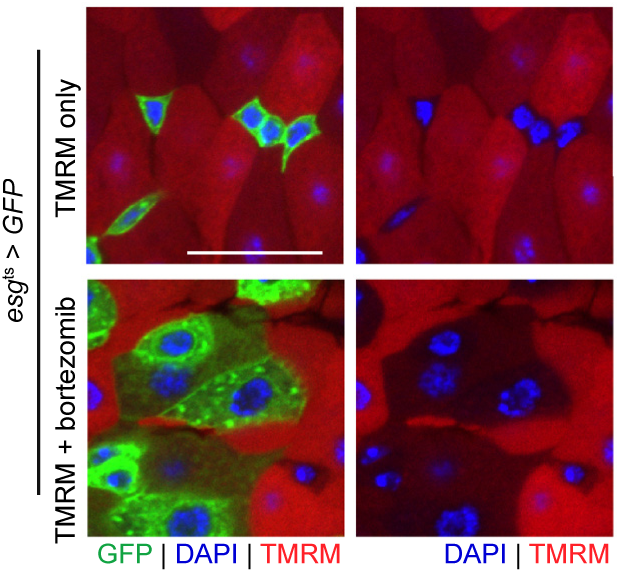
Intestinal stem/progenitor cells maintain lower TMRM accumulation than neighboring enterocytes during bortezomib treatment. Representative images of *esg*ᵗˢ > GFP intestines following 2 days of TMRM feeding alone or co-feeding with TMRM and bortezomib. GFP marks stem/progenitor cells (green), TMRM fluorescence is shown in red, and DAPI marks nuclei (blue). The GFP channel is omitted from the right-hand images to show relative TMRM fluorescence in the marked cells. Five guts were examined per condition.

**Supplementary Figure 6.**
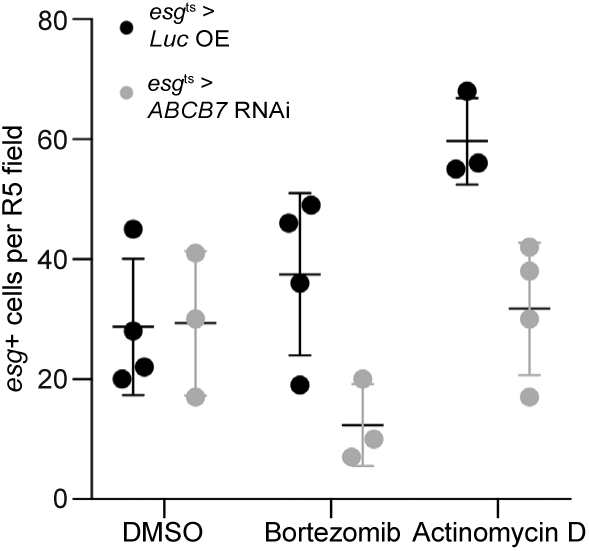
Quantification of *esg*+ stem/progenitor cells following *ABCB7* depletion during cytotoxic drug exposure. Adult flies expressing either *UAS-luciferase* (*Luc OE*) or *ABCB7* RNAi under *esg-GAL4*, *tub-GAL80^ts^* control were treated with DMSO, bortezomib, or actinomycin D. GFP-positive *esg*+ cells were counted within one imaged field of the R5 region per gut. Each point represents one gut; horizontal bars indicate mean and error bars indicate SD. DMSO: *n* = 4 *Luc OE* and 3 *ABCB7* RNAi; bortezomib: *n* = 4 *Luc OE* and 3 *ABCB7* RNAi; actinomycin D: *n* = 3 *Luc OE* and 4 *ABCB7* RNAi.

**Supplementary Figure 7.**
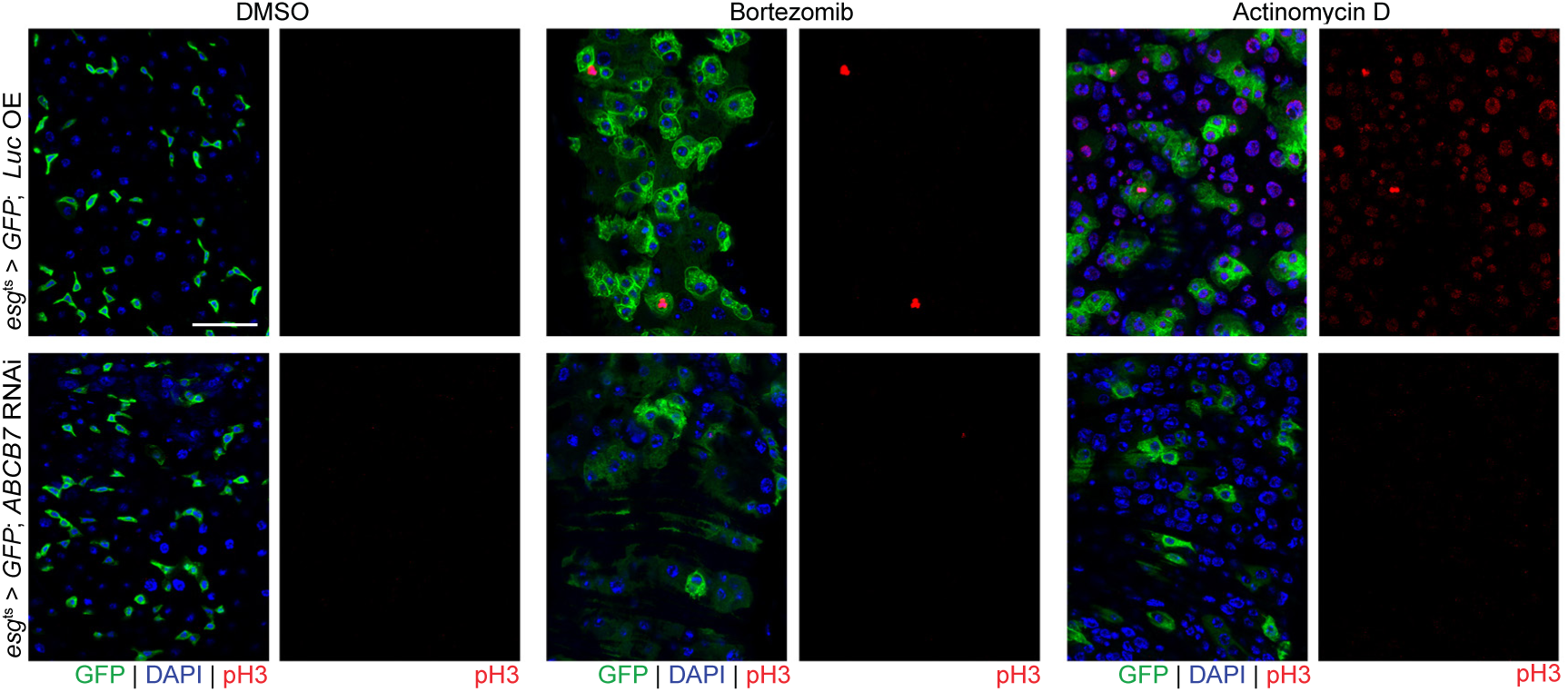
Representative pH3 staining following *ABCB7* knockdown. Representative images of control (*esg^ts^* > *Luc OE*) and *ABCB7* RNAi (*esg^ts^* > *ABCB7* RNAi) guts following DMSO, bortezomib, or actinomycin D treatment. DAPI is shown in blue, GFP in green, and pH3 in red. Scale bar, 30 µm.

**Supplementary Table I.**
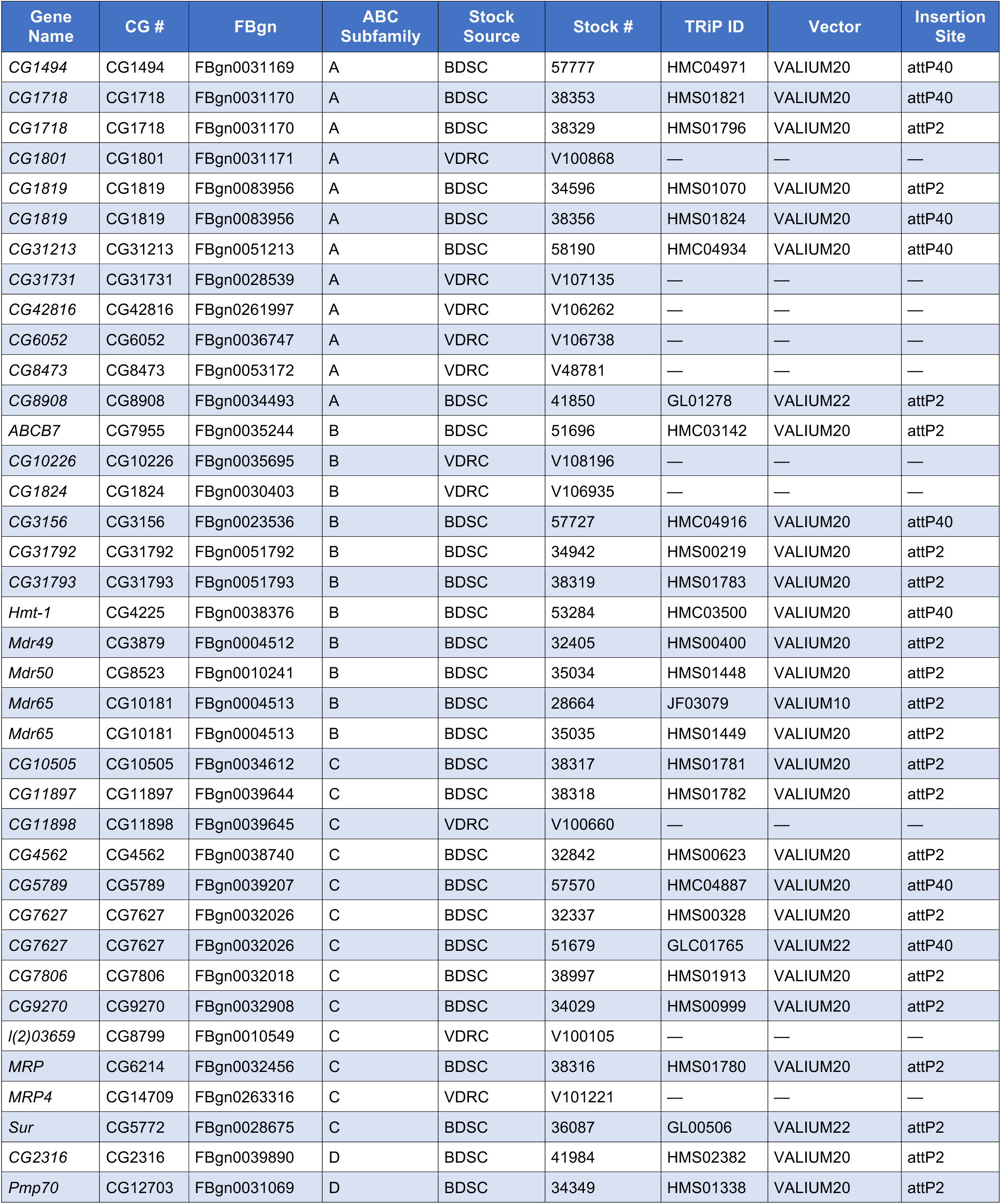
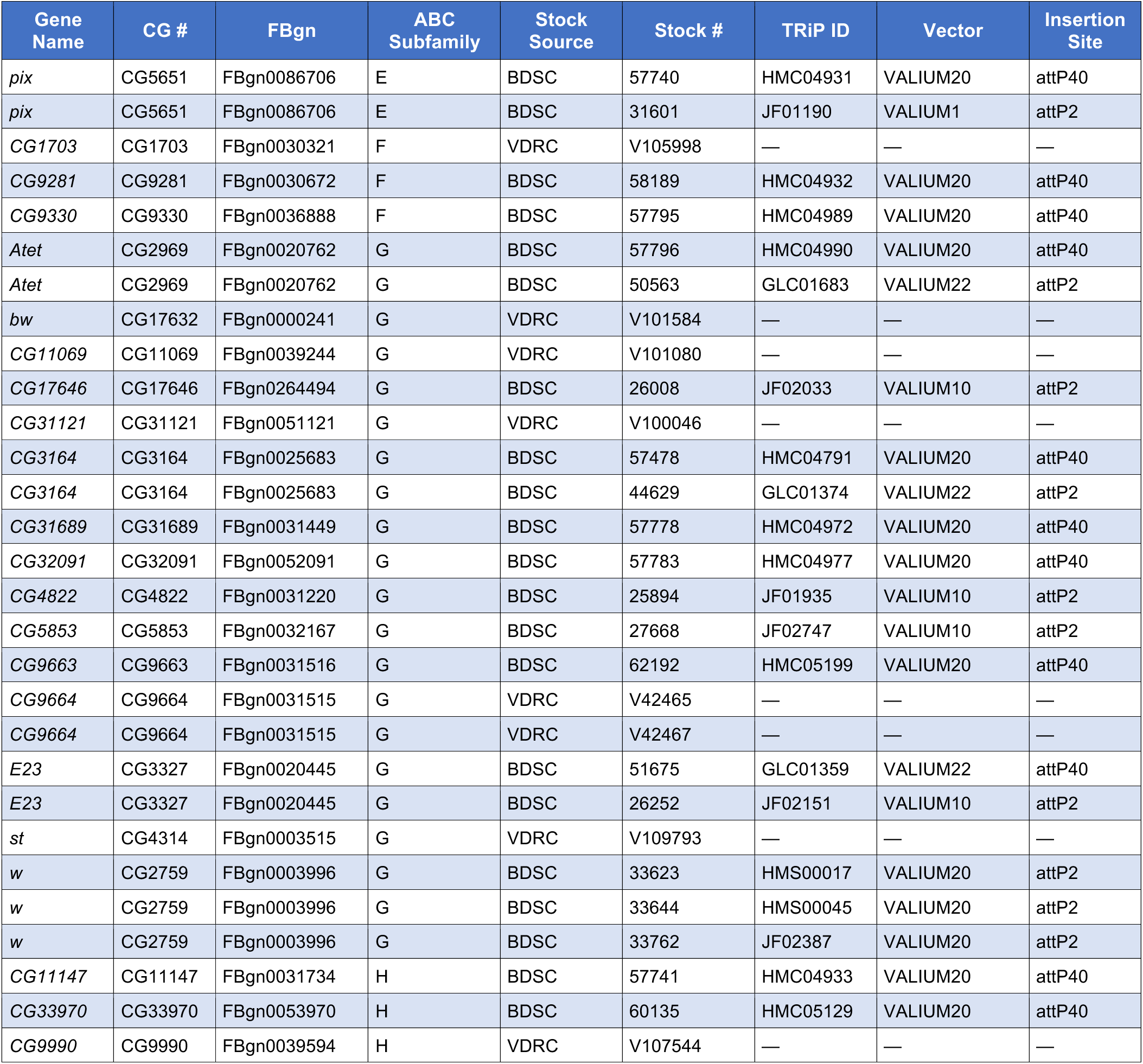
*Drosophila* ABC-family RNAi lines used in this study. *Complete list of UAS-RNAi transgenic lines targeting ABC transporter genes. Lines were obtained from the Bloomington Drosophila Stock Center (BDSC; TRiP collection) and the Vienna Drosophila Resource Center (VDRC). Gene names, CG identifiers, FlyBase gene numbers (FBgn), ABC subfamily classifications, stock numbers, TRiP line identifiers, RNAi vector backbones, and transgene insertion sites are indicated. Genes are sorted by ABC subfamily*.

## REFERENCES

1. Higgins, C. F. ABC transporters: from microorganisms to man. Annu. Rev. Cell Biol. 8, 67–113 (1992).

2. Locher, K. P., Lee, A. T. & Rees, D. C. The E. coli BtuCD structure: a framework for ABC transporter architecture and mechanism. Science 296, 1091–1098 (2002).

3. Kim, Y. & Chen, J. Molecular structure of human P-glycoprotein in the ATP-bound, outward-facing conformation. Science 359, 915–919 (2018).

4. Locher, K. P. Review. Structure and mechanism of ATP-binding cassette transporters. Philos. Trans. R. Soc. Lond. B Biol. Sci. 364, 239–245 (2009).

5. Wu, C., Chakrabarty, S., Jin, M., Liu, K. & Xiao, Y. Insect ATP-binding cassette (ABC) transporters: Roles in xenobiotic detoxification and Bt insecticidal activity. Int. J. Mol. Sci. 20, E2829 (2019).

6. Vasiliou, V., Vasiliou, K. & Nebert, D. W. Human ATP-binding cassette (ABC) transporter family. Hum. Genomics 3, 281–290 (2009).

7. Cho, Y. & Kim, Y. K. Cancer stem cells as a potential target to overcome multidrug resistance. Front. Oncol. 10, 764 (2020).

8. Robey, R. W. et al. Revisiting the role of ABC transporters in multidrug-resistant cancer. Nat. Rev. Cancer 18, 452–464 (2018).

9. Dean, M., Fojo, T. & Bates, S. Tumour stem cells and drug resistance. Nat. Rev. Cancer 5, 275–284 (2005).

10. Holohan, C., Van Schaeybroeck, S., Longley, D. B. & Johnston, P. G. Cancer drug resistance: an evolving paradigm. Nat. Rev. Cancer 13, 714–726 (2013).

11. Carnero, A. et al. The cancer stem-cell signaling network and resistance to therapy. Cancer Treat. Rev. 49, 25–36 (2016).

12. Muriithi, W. et al. ABC transporters and the hallmarks of cancer: roles in cancer aggressiveness beyond multidrug resistance. Cancer Biol. Med. 17, 253–269 (2020).

13. Goodell, M. A., Brose, K., Paradis, G., Conner, A. S. & Mulligan, R. C. Isolation and functional properties of murine hematopoietic stem cells that are replicating in vivo. J. Exp. Med. 183, 1797–1806 (1996).

14. Zhou, S. et al. The ABC transporter Bcrp1/ABCG2 is expressed in a wide variety of stem cells and is a molecular determinant of the side-population phenotype. Nat. Med. 7, 1028–1034 (2001).

15. Miguel-Aliaga, I., Jasper, H. & Lemaitre, B. Anatomy and Physiology of the Digestive Tract of Drosophila melanogaster. Genetics 210, 357–396 (2018).

16. Drozdzik, M., Czekawy, I., Oswald, S. & Drozdzik, A. Intestinal drug transporters in pathological states: an overview. Pharmacol. Rep. 72, 1173–1194 (2020).

17. Dietrich, C. G., Geier, A. & Oude Elferink, R. P. J. ABC of oral bioavailability: transporters as gatekeepers in the gut. Gut 52, 1788–1795 (2003).

18. Dekaney, C. M., Rodriguez, J. M., Graul, M. C. & Henning, S. J. Isolation and characterization of a putative intestinal stem cell fraction from mouse jejunum. Gastroenterology 129, 1567–1580 (2005).

19. Chee, Y. C. et al. Intrinsic xenobiotic resistance of the intestinal stem cell niche. Dev. Cell 46, 681–695.e5 (2018).

20. Micchelli, C. A. & Perrimon, N. Evidence that stem cells reside in the adult Drosophila midgut epithelium. Nature 439, 475–479 (2006).

21. Ohlstein, B. & Spradling, A. The adult Drosophila posterior midgut is maintained by pluripotent stem cells. Nature 439, 470–474 (2006).

22. Korzelius, J. et al. Escargot maintains stemness and suppresses differentiation in Drosophila intestinal stem cells. EMBO J. 33, 2967–2982 (2014).

23. Ohlstein, B. & Spradling, A. Multipotent Drosophila intestinal stem cells specify daughter cell fates by differential notch signaling. Science 315, 988–992 (2007).

24. Jiang, H. et al. Cytokine/Jak/Stat signaling mediates regeneration and homeostasis in the Drosophila midgut. Cell 137, 1343–1355 (2009).

25. Jiang, H., Tian, A. & Jiang, J. Intestinal stem cell response to injury: lessons from Drosophila. Cell. Mol. Life Sci. 73, 3337–3349 (2016).

26. Jiang, H. & Edgar, B. A. Intestinal stem cells in the adult Drosophila midgut. Exp. Cell Res. 317, 2780–2788 (2011).

27. Herrera, S. C. & Bach, E. A. JAK/STAT signaling in stem cells and regeneration: from Drosophila to vertebrates. Development 146, dev167643 (2019).

28. Mayer, F. et al. Evolutionary conservation of vertebrate blood-brain barrier chemoprotective mechanisms in Drosophila. J. Neurosci. 29, 3538–3550 (2009).

29. Denecke, S., Fusetto, R. & Batterham, P. Describing the role of Drosophila melanogaster ABC transporters in insecticide biology using CRISPR-Cas9 knockouts. Insect Biochem. Mol. Biol. 91, 1–9 (2017).

30. Denecke, S. et al. Characterization of a novel pesticide transporter and P-glycoprotein orthologues in Drosophila melanogaster. Proc. Biol. Sci. 289, 20220625 (2022).

31. Markstein, M. et al. Systematic screen of chemotherapeutics in Drosophila stem cell tumors. Proc. Natl. Acad. Sci. U. S. A. 111, 4530–4535 (2014).

32. Marianes, A. & Spradling, A. C. Physiological and stem cell compartmentalization within the Drosophila midgut. Elife 2, e00886 (2013).

33. Dutta, D. et al. Regional Cell-Specific Transcriptome Mapping Reveals Regulatory Complexity in the Adult Drosophila Midgut. Cell Rep. 12, 346–358 (2015).

34. Delacour, D., Salomon, J., Robine, S. & Louvard, D. Plasticity of the brush border — the yin and yang of intestinal homeostasis. Nat. Rev. Gastroenterol. Hepatol. 13, 161–174 (2016).

35. Antonello, Z. A., Reiff, T., Ballesta-Illan, E. & Dominguez, M. Robust intestinal homeostasis relies on cellular plasticity in enteroblasts mediated by miR-8-Escargot switch. EMBO J. 34, 2025–2041 (2015).

36. Morgan, N. S., Heintzelman, M. B. & Mooseker, M. S. Characterization of Myosin-IA and Myosin-IB, Two Unconventional Myosins Associated with theDrosophilaBrush Border Cytoskeleton. Dev. Biol. 172, 51–71 (1995).

37. Beebe, K., Lee, W.-C. & Micchelli, C. A. JAK/STAT signaling coordinates stem cell proliferation and multilineage differentiation in the Drosophila intestinal stem cell lineage. Dev. Biol. 338, 28–37 (2010).

38. Brand, A. H. & Perrimon, N. Targeted gene expression as a means of altering cell fates and generating dominant phenotypes. Development 118, 401–415 (1993).

39. McGuire, S. E., Mao, Z. & Davis, R. L. Spatiotemporal gene expression targeting with the TARGET and gene-switch systems in Drosophila. Sci. STKE 2004, l6 (2004).

40. Hölzl, H. et al. The regulatory complex of Drosophila melanogaster 26S proteasomes. Subunit composition and localization of a deubiquitylating enzyme. J. Cell Biol. 150, 119–130 (2000).

41. Lucchetta, E. M. & Ohlstein, B. The Drosophila midgut: a model for stem cell driven tissue regeneration. Wiley Interdiscip. Rev. Dev. Biol. 1, 781–788 (2012).

42. Scaduto, R. C., Jr & Grotyohann, L. W. Measurement of mitochondrial membrane potential using fluorescent rhodamine derivatives. Biophys. J. 76, 469–477 (1999).

43. Dermauw, W. & Van Leeuwen, T. The ABC gene family in arthropods: comparative genomics and role in insecticide transport and resistance. Insect Biochem. Mol. Biol. 45, 89–110 (2014).

44. Kondo, S. & Ueda, R. Highly improved gene targeting by germline-specific Cas9 expression in Drosophila. Genetics 195, 715–721 (2013).

45. Kondo, S. et al. Neurochemical organization of the Drosophila brain visualized by endogenously tagged neurotransmitter receptors. Cell Rep. 30, 284–297.e5 (2020).

46. Chen, Y.-C. D. et al. Using single-cell RNA sequencing to generate predictive cell-type-specific split-GAL4 reagents throughout development. Proc. Natl. Acad. Sci. U. S. A. 120, e2307451120 (2023).

47. Lee, P.-T. et al. A gene-specific T2A-GAL4 library for Drosophila. Elife 7, e35574 (2018).

48. Nagarkar-Jaiswal, S. et al. A library of MiMICs allows tagging of genes and reversible, spatial and temporal knockdown of proteins in Drosophila. Elife 4, (2015).

49. Regev, R., Katzir, H., Yeheskely-Hayon, D. & Eytan, G. D. Modulation of P-glycoprotein-mediated multidrug resistance by acceleration of passive drug permeation across the plasma membrane: MDR modulation by acceleration of drug permeation. FEBS J. 274, 6204–6214 (2007).

50. O’Connor, R. et al. The interaction of bortezomib with multidrug transporters: implications for therapeutic applications in advanced multiple myeloma and other neoplasias. Cancer Chemother. Pharmacol. 71, 1357–1368 (2013).

51. Zhou, S. et al. Bcrp1 gene expression is required for normal numbers of side population stem cells in mice, and confers relative protection to mitoxantrone in hematopoietic cells in vivo. Proc. Natl. Acad. Sci. U. S. A. 99, 12339–12344 (2002).

52. Wilson, B. J. et al. ABCB5 identifies a therapy-refractory tumor cell population in colorectal cancer patients. Cancer Res. 71, 5307–5316 (2011).

53. Schaedler, T. A. et al. Structures and functions of mitochondrial ABC transporters. Biochem. Soc. Trans. 43, 943–951 (2015).

54. Pondarre, C. et al. Abcb7, the gene responsible for X-linked sideroblastic anemia with ataxia, is essential for hematopoiesis. Blood 109, 3567–3569 (2007).

55. Lehrke, M. J. et al. The mitochondrial iron transporter ABCB7 is required for B cell development, proliferation, and class switch recombination in mice. Elife 10, e69621 (2021).

56. Tian, A. et al. Injury-stimulated Hedgehog signaling promotes regenerative proliferation of Drosophila intestinal stem cells. J. Cell Biol. 208, 807–819 (2015).

